# Enhanced Antimicrobial Potency and Therapeutic Index of a Defined C12 Quaternary Ammonium Compound

**DOI:** 10.64898/2026.10.29.762901

**Authors:** Madelyn M. Kist, I Gusti Ayu I. Wulandari, Joseph F. Buell, Lisa A. Morici, Chad J. Roy

## Abstract

Skin and soft tissue infections (SSTIs) remain a major clinical burden and are complicated by antimicrobial resistance, polymicrobial infections, and biofilm-mediated therapeutic tolerance. Topical antimicrobials can provide localized activity while reducing systemic exposure, but currently available agents are limited by narrow spectra, cytotoxicity, poor antibiofilm activity, and resistance development. We evaluated C12-alkyl(ethylbenzyl)dimethyl ammonium chloride (EQ12), a next-generation quaternary ammonium compound, *in vitro* as a topical antimicrobial candidate for leading bacterial causes of SSTIs. Broth microdilution assays were used to determine minimum inhibitory concentrations (MICs) and minimum bactericidal concentrations (MBCs) against methicillin-sensitive *Staphylococcus aureus* (MSSA), methicillin-resistant *S. aureus* (MRSA), *Acinetobacter baumannii*, *Klebsiella pneumoniae*, and *Pseudomonas aeruginosa*. Species-selective plating was used to assess bactericidal activity in staphylococcal-*P. aeruginosa* co-cultures. Crystal violet assays quantified biofilm inhibition against MRSA, *P. aeruginosa*, and *A. baumannii*, and serial passage experiments assessed resistance emergence over 14 passages. EQ12 inhibited all species tested, with MICs of 2.3 to 40 µg/mL and MBCs of 4.7 to 40 µg/mL. EQ12 outperformed benzalkonium chloride and povidone-iodine against Gram-negative isolates and retained bactericidal activity in mixed cultures, although staphylococcal susceptibility decreased in the presence of *P. aeruginosa*. EQ12 inhibited biofilm formation at sub-MIC to near-MIC concentrations and did not induce resistance in MRSA, *P. aeruginosa*, or *A. baumannii*, unlike comparator topical antibiotics. These findings support further development of EQ12 as a topical antimicrobial for SSTI-associated pathogens.

**Importance:** SSTIs are common, costly, and increasingly complicated by drug-resistant bacteria and biofilms. Topical antimicrobials are well suited for localized wound-associated infection, but many available agents are limited by toxicity, narrow spectrum, or resistance. This study shows that EQ12, a structurally defined C12-alkyl quaternary ammonium compound, has broad *in vitro* activity against clinically relevant Gram-positive and Gram-negative SSTI pathogens, inhibits biofilm formation, and presents a high barrier to rapid resistance development during repeated exposure. These findings identify EQ12 as a promising antimicrobial candidate and provide a bacteriological foundation for formulation and translational development.

## Introduction

Skin and soft tissue infections are among the most common reasons patients seek medical care in the United States, and their burden is likely underestimated since many cases are not culture confirmed (1, 2). Although Gram-positive organisms, particularly *S. aureus* and MRSA, dominate culture-confirmed SSTIs, Gram-negative pathogens including *P. aeruginosa*, *A. baumannii*, and *K. pneumoniae* are increasingly important in chronic, traumatic, burn, and health care-associated wound infections (2–5). Treatment is further complicated by antimicrobial resistance, biofilm-mediated tolerance, and polymicrobial community behavior that can alter antimicrobial susceptibility relative to monoculture conditions (6–9).

Topical antimicrobial therapy is often preferred for localized superficial or wound-associated infections because it can deliver high local concentrations with lower systemic exposure; however, current topical antibiotics and antiseptics have important limitations. Classical topical antibiotics can select for resistance and are often narrow in spectrum (10–13). Antiseptics can provide broad, multi-target activity, but their use may be constrained by cytotoxicity, formulation limitations, or reduced activity against some Gram-negative organisms and biofilms (14–17). These limitations create a need for topical agents that combine broad-spectrum bactericidal activity, antibiofilm activity, and a low propensity to select for resistance.

Quaternary ammonium compounds (QACs) are cationic membrane-active antimicrobials that disrupt bacterial membranes through physicochemical interactions rather than single-enzyme inhibition. Benzalkonium chloride (BZK), a widely used QAC, demonstrates broad activity but has limitations related to potency and cytotoxicity at practical concentrations (14, 18). The novel synthesized C12-alkyl(ethylbenzyl)dimethyl ammonium chloride, or colloquially ‘EQ12’, is a structurally defined QAC designed to retain membrane-disrupting antimicrobial activity while improving potency relative to conventional QAC comparators (**Figure 1**). The EQ12 molecule is conceptually related to (BZK), which is typically a mixture of alkyl-chain homologs, but the central structural distinction is that EQ12 is represented as a single C12-alkyl configuration. QAC activity depends on the cationic head group and hydrophobic alkyl chain: electrostatic interaction with negatively charged bacterial surfaces is followed by membrane insertion, disruption of membrane integrity, and cell death. Alkyl-chain length influences potency, membrane insertion, and concentration-dependent cytotoxicity, making the use of a defined C12 species a plausible strategy to tune antimicrobial effect while limiting host-cell damage.

**Figure 1.**
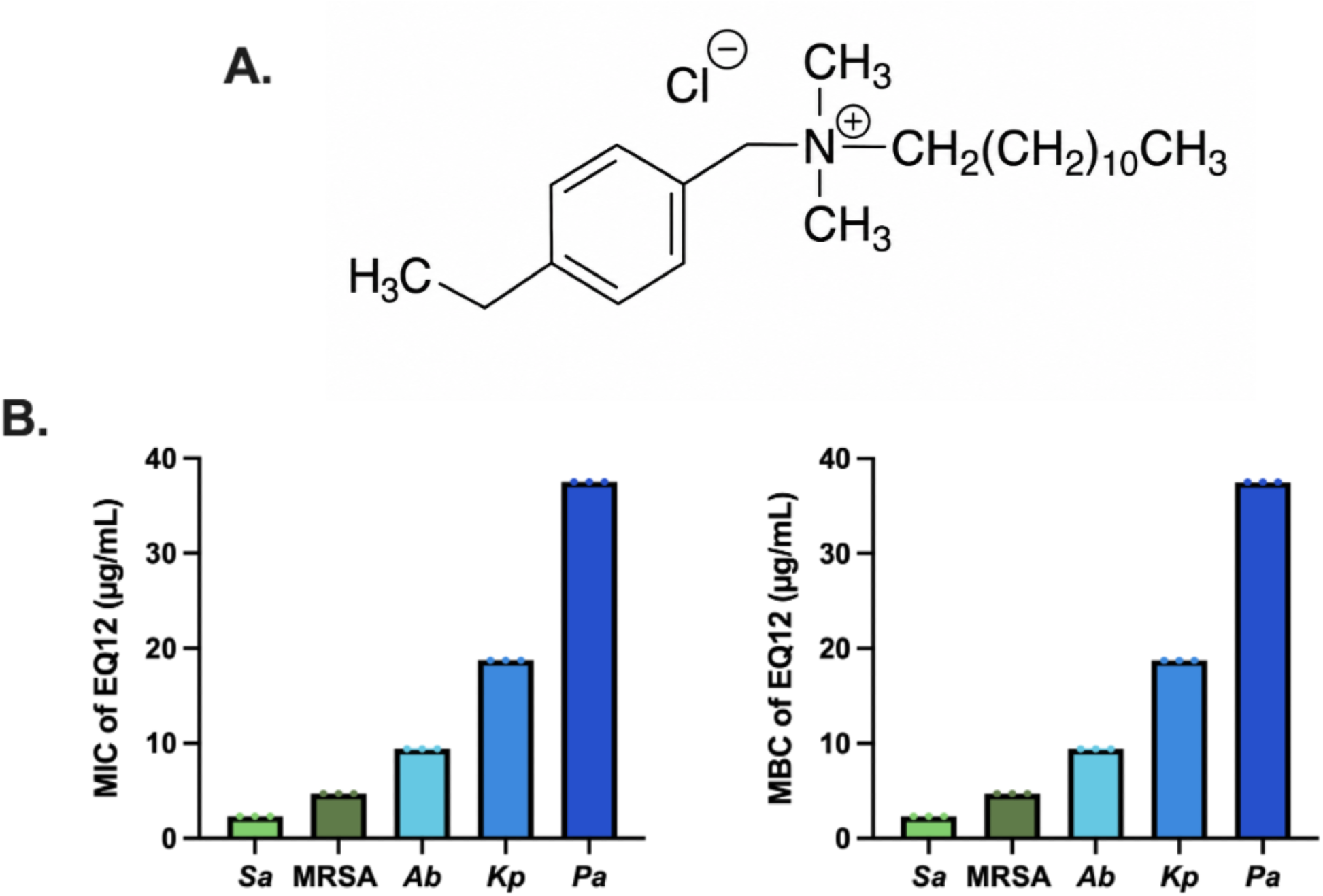
Chemical structure and *in vitro* antibacterial spectrum of EQ12. Panel A, chemical structure of C12-alkyl(ethylbenzyl)dimethyl ammonium chloride. Panel B, EQ12 MICs and MBCs against MSSA, MRSA, *A. baumannii*, *K. pneumoniae*, and *P. aeruginosa* determined by broth microdilution.

Here, we evaluated EQ12 using an *in vitro* bacteriological framework relevant to SSTI-associated pathogens. We tested bacteriostatic and bactericidal activity against individual bacterial species, bactericidal activity in *S. aureus*-*P. aeruginosa* and MRSA-*P. aeruginosa* co-cultures, inhibition of biofilm formation by MRSA, *P. aeruginosa*, and *A. baumannii*, and resistance emergence during repeated subinhibitory exposure.

## Materials and Methods

### Antimicrobial agents

Commercial antiseptics included BZK 50% w/w (CAS 63449-41-2; Thermo Fisher Scientific), chlorhexidine gluconate 2% (CHG; DurVet), and povidone-iodine 10% (Betadine; BD; Dynarex). These were diluted to assay-specific working concentrations in 0.22-micrometer filter-sterilized deionized water and stored in 1-mL aliquots at room temperature. EQ12 (C12-alkyl[ethylbenzyl]dimethyl ammonium chloride; CAS 14352-42-9; Bio Protectant Technologies) was prepared from powder as a 10-mg/mL stock solution in 0.22-micrometer filter-sterilized deionized water and stored in 1-mL aliquots at room temperature. Gentamicin (GEN; CAS 1405-41-0; Thermo Fisher Scientific) and bacitracin (BAC; CAS 1405-87-4; Thermo Fisher Scientific) were dissolved in deionized water, filter sterilized, and stored in 1-mL aliquots at -20 degrees C. Mupirocin (MUP; CAS 12650-69-0; TCI America) was dissolved in 10% aqueous ethanol, heated for approximately 5 min at 50 to 55 degrees C to facilitate dissolution, filter sterilized, and stored in 1-mL aliquots at -20 degrees C. Antimicrobial stocks were diluted into assay-specific media immediately before use.

### Bacterial strains and culture conditions

Antimicrobial activity was evaluated against MSSA, MRSA, *A. baumannii*, *K. pneumoniae*, and *P. aeruginosa*. From freshly streaked LB agar plates, three colonies of each species were inoculated separately into 5 mL Luria-Bertani broth and incubated for 16 to 18 h at 37 degrees C with shaking at 200 rpm. Overnight cultures were diluted 1:100 in fresh LB broth and grown to an optical density at 600 nm of 0.75 before dilution to 5 x 10^5 CFU/mL in cation-adjusted Mueller-Hinton II broth.

### MIC and MBC assays

MICs were determined by broth microdilution. Bacterial suspensions were added to round-bottom tissue culture-treated 96-well plates containing 2-fold serial dilutions of EQ12 or comparators in Mueller-Hinton broth. Plates were sealed and incubated for 18 h at 37 degrees C with shaking at 200 rpm. Each plate included sterility controls containing medium and antimicrobial without bacteria. The MIC was defined as the lowest antimicrobial concentration at which no turbidity was observed. For MBC determination, 10 microliters from each well was spot plated onto LB agar and incubated statically for 24 h at 37 degrees C. The MBC was defined as the lowest concentration with no visible growth. Values represent means from three independent biological replicates.

### Mixed-species bactericidal assays

To evaluate whether bactericidal activity changed in a simulated co-infection environment, MSSA-*P. aeruginosa* and MRSA-*P. aeruginosa* co-cultures were prepared at 5 x 10^5 CFU/mL of each species, yielding a total bacterial density of 1 x 10^6 CFU/mL. Co-cultures were incubated with 2-fold serial dilutions of antimicrobial agents in Mueller-Hinton broth for 18 h at 37 degrees C with shaking. To determine species-specific MBCs after co-culture exposure, 10 microliters from each well was spot plated onto mannitol salt agar for *Staphylococcal* isolation and *Pseudomonas* isolation agar for *P. aeruginosa* isolation. Plates were incubated statically for 24 h at 37 degrees C. Co-culture MBCs were compared with corresponding monoculture MBCs.

### Biofilm inhibition assays

Biofilm inhibition was assessed by adapting static crystal violet assays (19, 20). For MRSA, overnight cultures were prepared in brain heart infusion broth supplemented with 1% glucose. For *P. aeruginosa* and *A. baumannii*, overnight cultures were prepared in tryptic soy broth supplemented with 1% glucose. Cultures were diluted in fresh species-specific medium, added to flat-bottom untreated 96-well plates containing 2-fold serial dilutions of antimicrobial agents, and incubated statically for exactly 24 h at 37 degrees C to allow biofilm formation in the presence of antimicrobials. Wells were gently aspirated, washed three times with PBS, stained with 0.1% crystal violet for 15 min, washed three times with deionized water, and solubilized with 100% ethanol. Absorbance was measured at 570 nm. Background values from medium-antimicrobial wells without bacteria were subtracted. The minimum biofilm inhibition concentration (MBIC) was defined as the lowest concentration at which OD570 was less than 0.1. Percent biofilm inhibition was calculated as |1 - (treated/control)| x 100, where control represented untreated mature biofilm wells. MBIC-to-MIC conversion was calculated as mean MBIC divided by mean MIC.

### Serial passage resistance assays

Serial passage experiments were performed independently for MRSA, *P. aeruginosa*, and *A. baumannii* over 28 days. Initial MICs were determined by broth microdilution. Bacteria from the well containing the highest antimicrobial concentration that still permitted visible growth, corresponding to one dilution below the MIC, were diluted 1:100 into fresh LB broth, incubated overnight, and used for the next broth microdilution assay. This process was repeated for 14 passages. Resistance was defined as a greater than 4-fold increase in MIC relative to the initial passage.

### Statistical analysis

Statistical analyses and figure generation were performed using GraphPad Prism 10.3.1. Ordinary one-way analysis of variance with multiple comparisons was used to compare EQ12 with comparator agents for single-species MIC, MBC, and biofilm data. Welch’s t test was used to compare co-culture MBCs with corresponding single-species MBCs. *P* values below 0.05 were considered significant.

## Results

EQ12 exhibits broad-spectrum activity against SSTI-associated pathogens. EQ12 inhibited the growth of all five species tested, with MICs ranging from 2.3 to 40 µg/mL and MBCs ranging from 4.7 to 40 µg/mL (**Table 1**, **Figure 1**). Activity was strongest against staphylococci and lowest against *P. aeruginosa*, consistent with a Gram-positive-skewed but broad-spectrum profile. EQ12 was most potent against MSSA (MIC, 2.3 µg/mL; MBC, 4.7 µg/mL) and least potent against *P. aeruginosa* (MIC and MBC, 40 µg/mL). This species-dependent trend was similar to that observed for BZK but with greater overall potency.

**Table 1A.** MIC values (µg/mL) for EQ12 and comparators against SSTI-associated pathogens.

| Species | EQ12 | BZK | CHG | BD | BAC | MUP | GEN |
| --- | --- | --- | --- | --- | --- | --- | --- |
| MSSA | 2.3 | 3.125 | 3.125 | 1000 | 32.5 | 0.125 | - |
| MRSA | 4.7 | 6.25 | 3.125 | 500 | 37.5 | 0.125 | - |
| *A. baumannii* | 9.4 | 25 | 12.5 | 2000 | - | - | 3.1 |
| *K. pneumoniae* | 18.8 | 25 | 12.5 | 4000 | - | - | 1.56 |
| *P. aeruginosa* | 40 | 100 | 12.5 | 4000 | - | - | 1 |

**Table 1B.** MBC values (µg/mL) for EQ12 and comparators against SSTI-associated pathogens.

| Species | EQ12 | BZK | CHG | BD | BAC | MUP | GEN |
| --- | --- | --- | --- | --- | --- | --- | --- |
| MSSA | 4.7 | 16.7 | 20.8 | 4000 | 21.7 | - | - |
| MRSA | 4.7 | 10.4 | 3.125 | 500 | 37.5 | - | - |
| *A. baumannii* | 9.4 | 25 | 12.5 | 2000 | - | - | 3.1 |
| *K. pneumoniae* | 18.8 | 50 | 41.7 | 4000 | - | - | 1.56 |
| *P. aeruginosa* | 40 | 100 | 12.5 | 4000 | - | - | 1 |

Against MSSA, EQ12 was more bacteriostatic than all comparators except mupirocin, and more bactericidal than bacitracin and povidone-iodine (**Figure 2**). Against MRSA, EQ12 was more bacteriostatic and more bactericidal than BZK, povidone-iodine, and bacitracin. Against *A. baumannii*, EQ12 was more bacteriostatic and bactericidal than BZK, CHG, and povidone-iodine, although gentamicin was more potent. Against *K. pneumoniae*, EQ12 was more bacteriostatic than BZK and povidone-iodine and more bactericidal than BZK, CHG, and povidone-iodine. Against *P. aeruginosa*, EQ12 outperformed BZK and povidone-iodine but was less potent than CHG and gentamicin. Co-culture with *P. aeruginosa* reduces staphylococcal susceptibility to EQ12 but does not reduce *P. aeruginosa* susceptibility.

**Figure 2.**
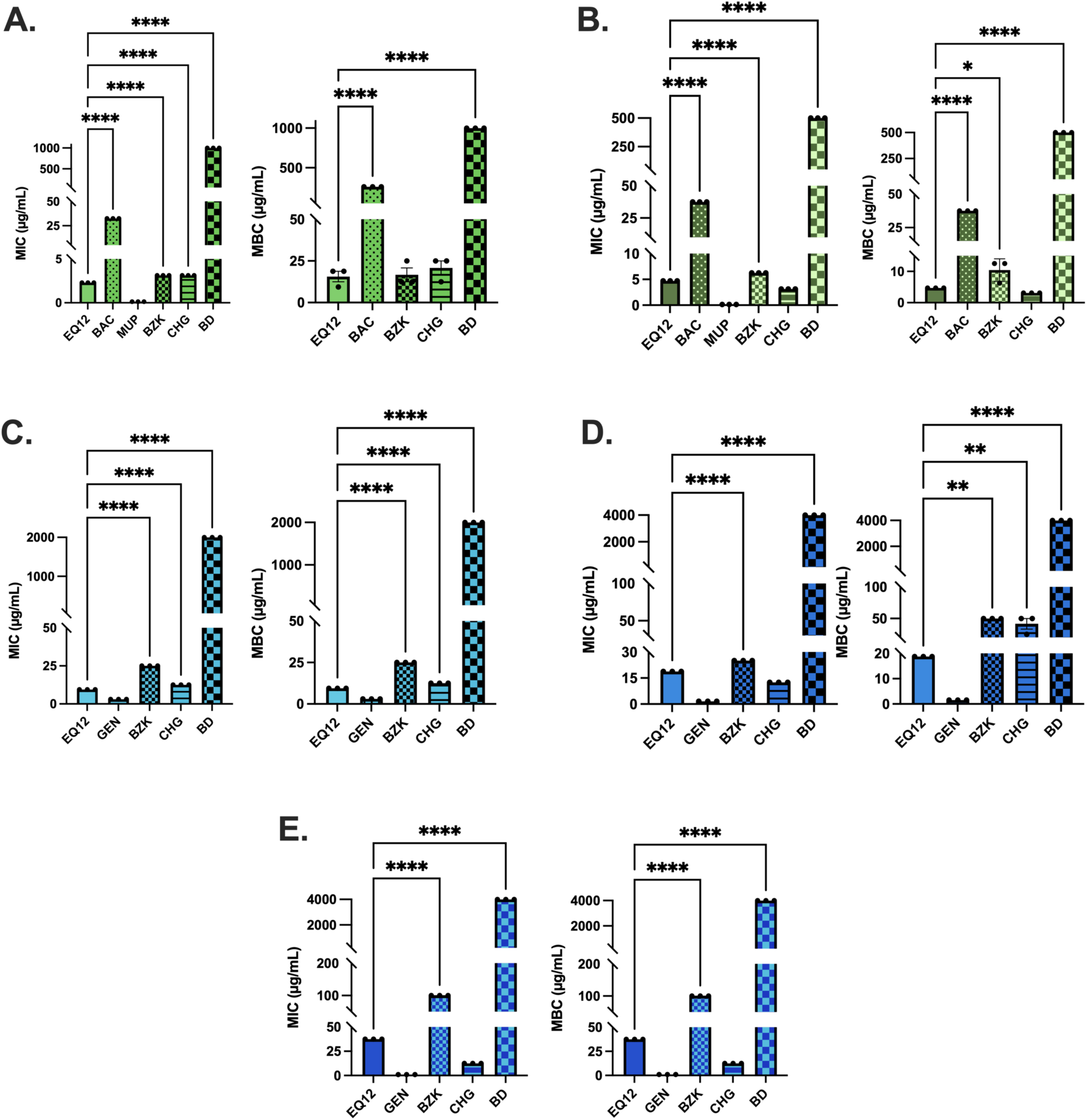
Comparative antibacterial activity of EQ12 against Gram-positive and Gram-negative SSTI pathogens SSTI pathogens. MIC and MBC values for EQ12 and comparator topical antimicrobials against MSSA, MRSA, *A. baumannii*, *K. pneumoniae*, and *P. aeruginosa*. Statistical comparisons were performed using ordinary one-way ANOVA with multiple comparisons.

Because wound-associated SSTIs can be polymicrobial, we next compared bactericidal activity in staphylococcal-*P. aeruginosa* co-cultures. In MRSA-*P. aeruginosa* co-culture, the EQ12 MBC against MRSA increased from 4.7 to 37.5 µg/mL, whereas the EQ12 MBC against *P. aeruginosa* remained similar to monoculture values (37.5 to 50 µg/mL) (**Table 2**). In MSSA-*P. aeruginosa* co-culture, the EQ12 MBC against MSSA similarly increased from 4.7 to 37.5 µg/mL, while the *P. aeruginosa* MBC remained unchanged at 37.5 µg/mL (**Table 2**). BZK showed a similar trend but required higher concentrations than EQ12. CHG showed no statistically significant monoculture-to-co-culture shifts, whereas povidone-iodine and bacitracin showed increased staphylococcal MBCs in co-culture. Gentamicin MBCs against *P. aeruginosa* increased from 1 to 2 µg/mL in the presence of either staphylococcal species. These data indicate that the presence of *P. aeruginosa* reduces staphylococcal susceptibility to several agents, including EQ12, but does not meaningfully reduce *P. aeruginosa* susceptibility to EQ12 (**Figure 3**).

**Figure 3.**
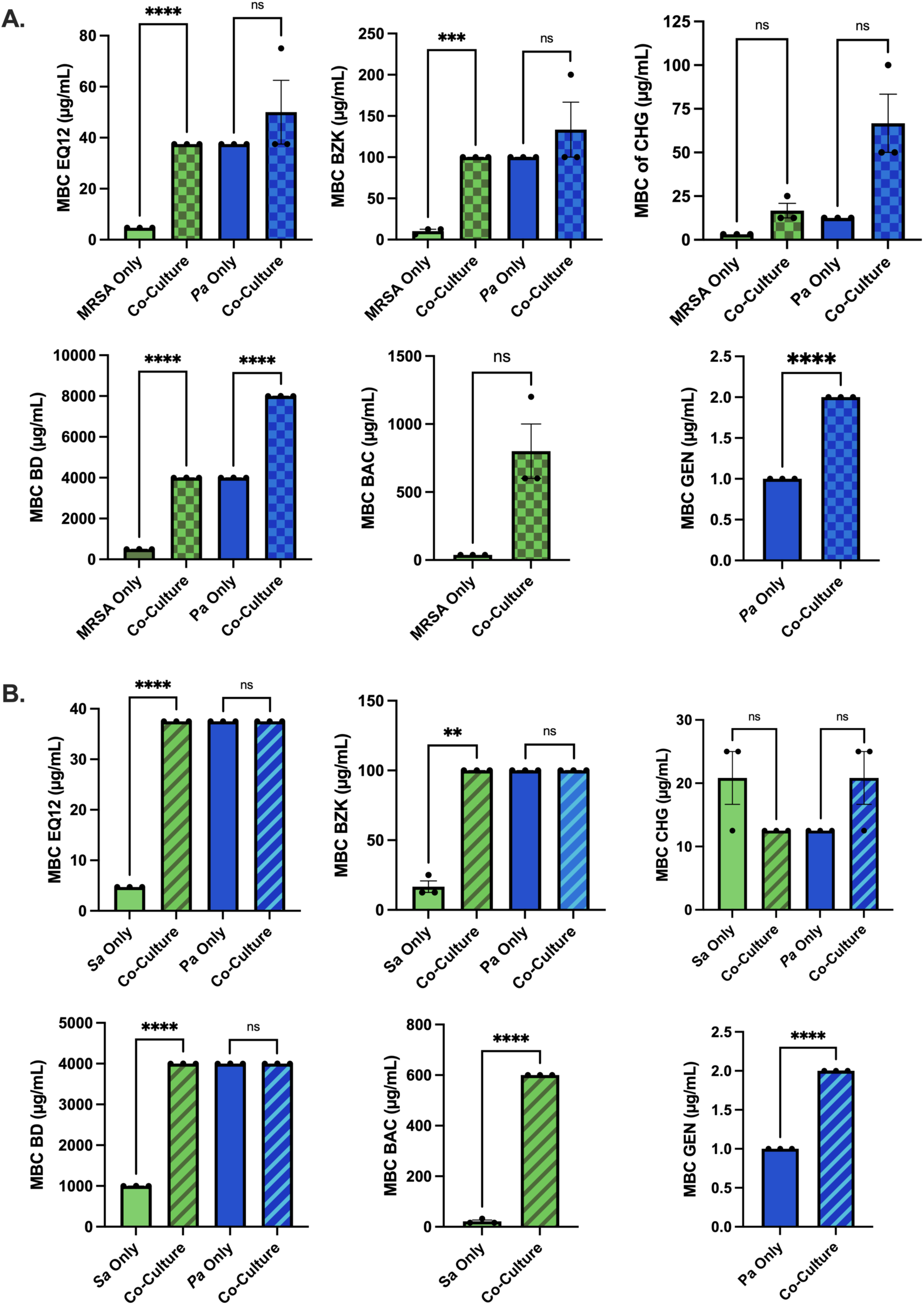
Bactericidal activity of EQ12 in *Staphylococcal-P. aeruginosa* co-cultures. MRSA-*P. aeruginosa* and MSSA-*P. aeruginosa* co-cultures were treated with EQ12 or comparator agents, and species-specific MBCs were determined by plating on mannitol salt agar or *Pseudomonas* isolation agar.

**Table 2.** Selected MBC shifts in *staphylococcal*-*P. aeruginosa* co-cultures.

| Co-culture | Species recovered | Agent | Alone MBC | Co-culture MBC | Significant change |
| --- | --- | --- | --- | --- | --- |
| MRSA-*P. aeruginosa* | MRSA | EQ12 | 4.7 | 37.5 | Yes |
| MRSA-*P. aeruginosa* | *P. aeruginosa* | EQ12 | 37.5 | 50 | No |
| MSSA-*P. aeruginosa* | MSSA | EQ12 | 4.7 | 37.5 | Yes |
| MSSA-*P. aeruginosa* | *P. aeruginosa* | EQ12 | 37.5 | 37.5 | No |
| MRSA-*P. aeruginosa* | MRSA | BZK | 10.4 | 100 | Yes |
| MRSA-*P. aeruginosa* | *P. aeruginosa* | BZK | 100 | 133.3 | No |
| MSSA-*P. aeruginosa* | MSSA | BZK | 10.4 | 100 | Yes |
| MSSA-*P. aeruginosa* | *P. aeruginosa* | BZK | 100 | 100 | No |
| MRSA-*P. aeruginosa* | MRSA | BD | 500 | 4000 | Yes |
| MRSA-*P. aeruginosa* | *P. aeruginosa* | BD | 4000 | 8000 | Yes |
| MSSA-*P. aeruginosa* | MSSA | BD | 1000 | 4000 | Yes |
| MSSA-*P. aeruginosa* | *P. aeruginosa* | BD | 4000 | 4000 | No |
| MRSA-*P. aeruginosa* | MRSA | BAC | 37.5 | 800 | No |
| MSSA-*P. aeruginosa* | MSSA | BAC | 21.7 | 600 | Yes |
| MRSA-*P. aeruginosa* | *P. aeruginosa* | GEN | 1 | 2 | Yes |
| MSSA-*P. aeruginosa* | *P. aeruginosa* | GEN | 1 | 2 | Yes |

EQ12 inhibits MRSA, *P. aeruginosa*, and *A. baumannii* biofilm formation. EQ12 inhibited biofilm formation by all tested species at sub-MIC to near-MIC concentrations (**Table 3**, **Figure 4**). The EQ12 MBIC was 2.3 µg/mL against MRSA, 37.5 µg/mL against *P. aeruginosa*, and 4.7 µg/mL against *A. baumannii*, corresponding to 0.5x, 0.9x, and 0.5x the respective MICs. BZK showed similar fold-MIC MBIC values but required higher absolute concentrations for *P. aeruginosa* and *A. baumannii*. At MIC concentrations, EQ12 inhibited biofilm formation by 99.1% for MRSA, 96.2% for *P. aeruginosa*, and 103.1% for *A. baumannii*. Across antiseptic comparators, EQ12 produced the highest percent inhibition values, with BZK ranging from 95.8 to 99.4%, CHG from 76.3 to 95.4%, and povidone-iodine from 76.8 to 93.2%. Gentamicin showed comparatively weaker biofilm inhibition against *P. aeruginosa* and *A. baumannii* at its MIC. These data support EQ12 activity against planktonic organisms and early biofilm formation.

**Figure 4.**
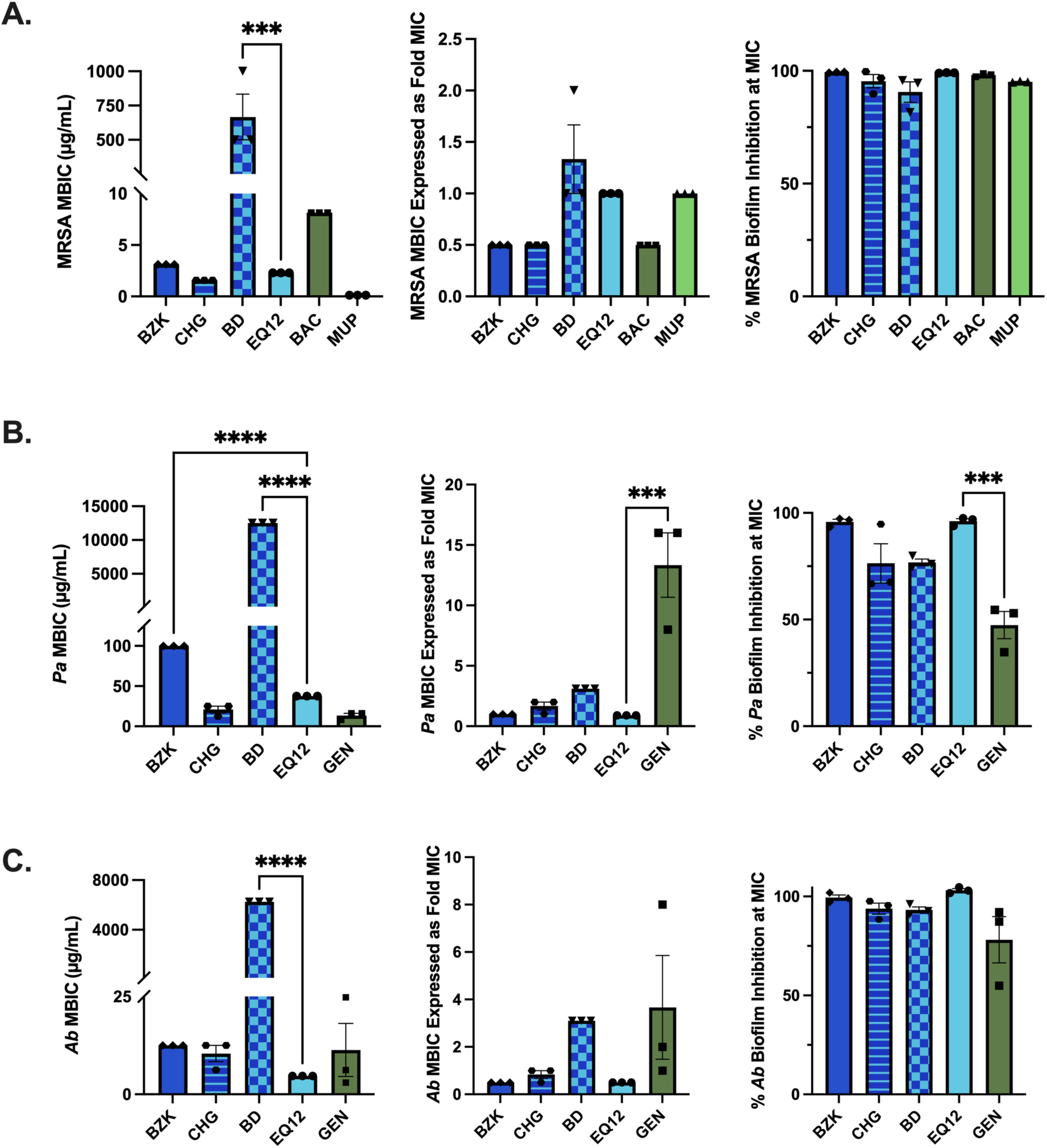
EQ12 inhibits biofilm formation by MRSA, *P. aeruginosa*, and *A. baumannii*. Biofilms were formed for 24 h in the presence of EQ12 or comparators, stained with crystal violet, solubilized with ethanol, and read at OD570. MBICs, fold-MIC values, and percent inhibition at MIC are shown for A. MRSA, B. *P. aeruginosa*, and C. *A. baumannii*.

**Table 3.** Biofilm inhibition by EQ12 and comparator agents.

| Species | Agent | MBIC (micrograms/mL) | MBIC/MIC | Biofilm inhibition at MIC (%) |
| --- | --- | --- | --- | --- |
| MRSA | EQ12 | 2.3 | 0.5x | 99.1 |
| MRSA | BZK | 3.125 | 0.5x | 99.4 |
| MRSA | CHG | 1.56 | 0.5x | 95.4 |
| MRSA | BD | 666.7 | 1.3x | 90.6 |
| MRSA | BAC | 8.15 | 0.2x | 98.1 |
| MRSA | MUP | 0.125 | 1x | 95.2 |
| *P. aeruginosa* | EQ12 | 37.5 | 0.9x | 96.2 |
| *P. aeruginosa* | BZK | 100 | 1x | 95.8 |
| *P. aeruginosa* | CHG | 20.8 | 1.6x | 76.3 |
| *P. aeruginosa* | BD | 12500 | 3.1x | 76.8 |
| *P. aeruginosa* | GEN | 13.3 | 6.6x | 47.4 |
| *A. baumannii* | EQ12 | 4.7 | 0.5x | 103.1 |
| *A. baumannii* | BZK | 12.5 | 0.5x | 99.4 |
| *A. baumannii* | CHG | 10.4 | 0.8x | 93.8 |
| *A. baumannii* | BD | 6250 | 3.1x | 93.2 |
| *A. baumannii* | GEN | 11.3 | 3.6x | 78.1 |

EQ12 does not induce rapid resistance under repeated *in vitro* exposure (**Figure 5**). Serial passage assays were used to determine whether repeated exposure at one dilution below the MIC selected for reduced susceptibility. None of the three bacterial species developed resistance to EQ12, BZK, povidone-iodine, or bacitracin over 14 passages (**Table 4**). EQ12 MICs remained completely stable against MRSA and *P. aeruginosa* and varied only modestly against *A. baumannii*, remaining below the resistance threshold. In contrast, resistance emerged rapidly to classical topical antibiotic comparators. MRSA developed mupirocin resistance at passage 8, reaching a 64-fold MIC increase by passage 14. *P. aeruginosa* developed gentamicin resistance at passage 5, reaching a 64-fold MIC increase by passage 14. *A. baumannii* developed gentamicin resistance at passage 3, reaching a 512-fold MIC increase by passage 14. CHG produced a transient resistance-threshold crossing in *P. aeruginosa* at passage 5, but the MIC subsequently decreased and remained stable below the resistance threshold. These findings indicate that EQ12 presents a high barrier to rapid resistance acquisition under the conditions tested.

**Figure 5.**
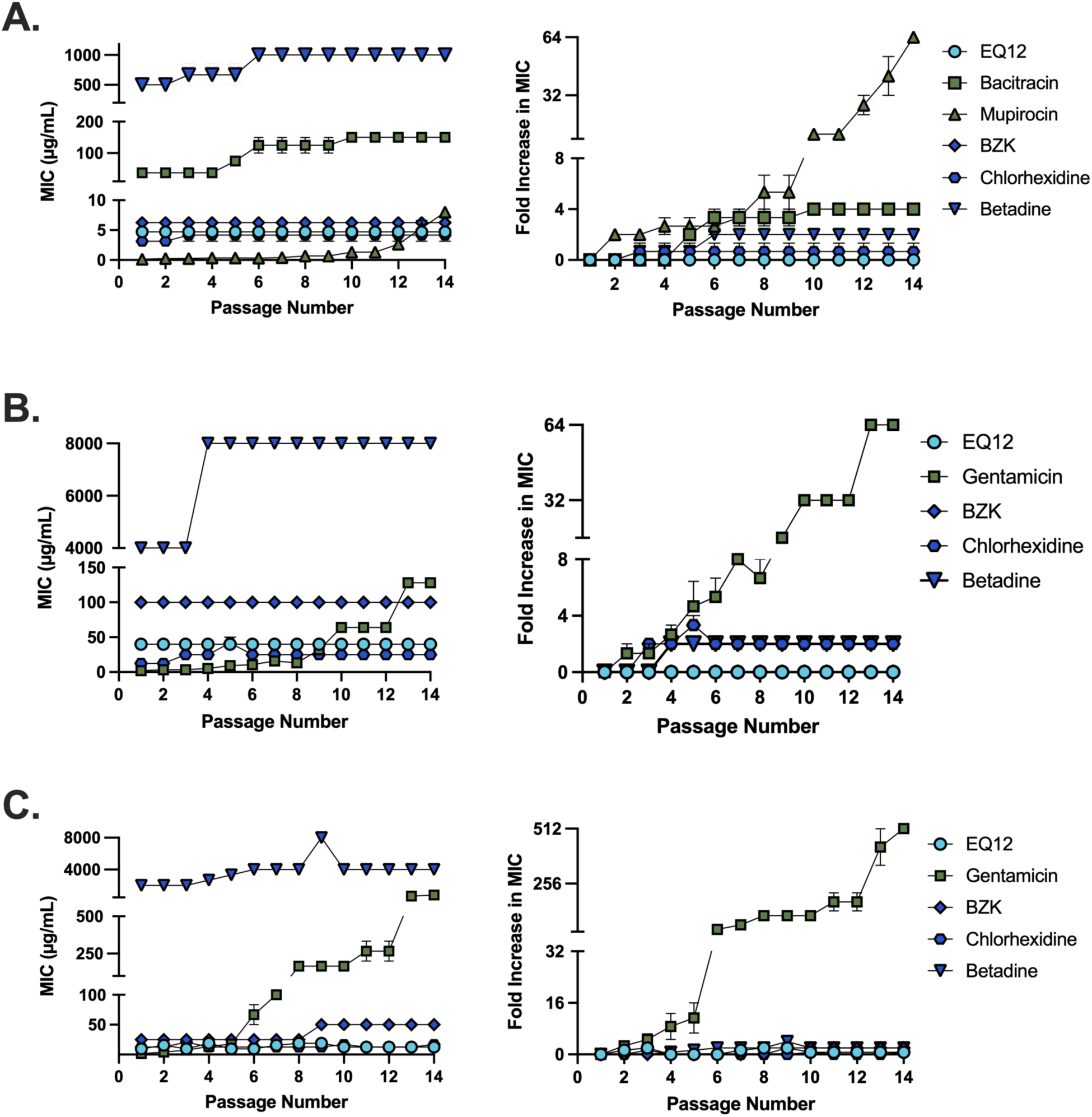
Repeated exposure to EQ12 does not induce rapid resistance in MRSA, *P. aeruginosa*, or *A. baumannii*. Bacteria were serially passaged for 14 cycles in EQ12 or comparator agents. MICs were for determined after each passage for A. MRSA, B. *P. aeruginosa*, or C. *A. baumannii*. Resistance was defined as a greater than 4-fold increase relative to the initial MIC.

**Table 4.** Serial-passage resistance summary over 14 passages.

| Species | Agent with resistance emergence | Passage at threshold crossing | Final fold MIC increase | EQ12 resistance observed |
| --- | --- | --- | --- | --- |
| MRSA | Mupirocin | 8 | 64-fold | No |
| *P. aeruginosa* | Gentamicin | 5 | 64-fold | No |
| *A. baumannii* | Gentamicin | 3 | 512-fold | No |

## Discussion

Synthesis of EQ12 was motivated by the hypothesis that enriching the structurally defined C12-alkyl component could enhance antimicrobial potency and reduce the concentrations of quaternary ammonium compounds required for human health applications. Accordingly, this study provides a focused *in vitro* comparative evaluation of EQ12 as a candidate topical antimicrobial for SSTI-relevant bacteria. EQ12 demonstrated bacteriostatic and bactericidal activity against both Gram-positive and Gram-negative pathogens, inhibited biofilm formation by MRSA, *P. aeruginosa*, and *A. baumannii*, and did not select for resistance during 14 serial passages. The aggregate profile differentiates EQ12 from traditional topical antibiotics which, although potent, showed narrower activity and rapid resistance emergence in the serial passage assays.

EQ12 potency was greatest against staphylococci and lowest against *P. aeruginosa*, reflecting the well-established increased intrinsic tolerance of Gram-negative bacteria to many membrane-active compounds. The outer membrane, efflux systems, and adaptive envelope responses of Gram-negative organisms can reduce susceptibility to cationic antimicrobials, particularly in *P. aeruginosa* and *A. baumannii* (4, 21–23). Despite this expected trend, EQ12 remained bactericidal against all tested Gram-negative species and outperformed BZK and povidone-iodine against both *P. aeruginosa* and *A. baumannii*. This is notable because BZK is a closely related QAC comparator, suggesting that EQ12 structure and alkyl-chain composition may translate into improved absolute potency while retaining a membrane-active mechanism.

The co-culture data emphasize the importance of testing topical antimicrobials under conditions that begin to approximate polymicrobial SSTIs. In both MRSA-*P. aeruginosa* and MSSA-*P. aeruginosa* co-cultures, the EQ12 MBC against staphylococci increased, whereas the MBC against *P. aeruginosa* was preserved. The same directional pattern was observed for BZK, and larger staphylococcal shifts occurred with povidone-iodine and bacitracin. These data suggest that *P. aeruginosa* can alter staphylococcal susceptibility in mixed cultures, potentially through community-level tolerance, metabolic interactions, matrix-associated protection, or differential survival after antimicrobial exposure. Because the present assay used a simplified planktonic co-culture model and species-selective plating, future studies should define the mechanism responsible for the staphylococcal shift and test more complex polymicrobial biofilm models.

Biofilm formation contributes to recalcitrant wound infection and can reduce antimicrobial efficacy through matrix sequestration, altered physiology, and heterogeneous growth states (7, 19, 24). EQ12 inhibited biofilm formation at concentrations at or below the MIC for all tested biofilm-forming species. Importantly, EQ12 showed strong activity against *P. aeruginosa* and *A. baumannii*, two organisms for which biofilm formation is central to persistence in wounds and medical environments (4, 5, 21, 25). These assays measured inhibition of biofilm formation during antimicrobial exposure rather than eradication of established mature biofilms; therefore, the results support a role for EQ12 in preventing or limiting biofilm development but should not be overinterpreted as mature biofilm eradication.

The serial passage experiments provide an important comparative signal. As expected, mupirocin and gentamicin selected for resistance rapidly under repeated subinhibitory exposure. In contrast, EQ12 did not select for resistance in MRSA, *P. aeruginosa*, or *A. baumannii* over 14 passages. This pattern is consistent with the multi-target membrane-active mechanism proposed for QACs and other antiseptics, which can increase the barrier to target-specific resistance relative to classical antibiotics (14).

Nonetheless, QAC tolerance and efflux-associated reductions in susceptibility have been reported for some bacteria, and the absence of resistance in this study should be interpreted within the tested conditions and timeframe. Longer passaging, genomic analysis, efflux-pump profiling, and cross-tolerance studies will be important for defining the durability of the EQ12 resistance barrier.

This study has limitations, including assays using reference-style *in vitro* conditions that do not reproduce the full biochemical complexity of wound exudate, host proteins, necrotic tissue, pH variation, or mature polymicrobial biofilms. Cytocompatibility, formulation stability, tissue compatibility, and activity in protein-rich wound-like matrices also require further testing. The data promote a strong microbiological rationale for continued development of EQ12 as a topical antimicrobial candidate. EQ12 demonstrates broad-spectrum *in vitro* antibacterial activity, retained activity in simplified mixed-species co-cultures, inhibited biofilm formation, and did not induce rapid resistance under repeated exposure. These properties support further nonclinical development of EQ12 for topical management of SSTI-associated bacterial contamination and wound-associated microbial burden.

## Acknowledgments

We thank Bio Protectant Technologies, Inc., for collaboration and for providing EQ12. Figures were created with BioRender, and statistical analyses were conducted in GraphPad Prism.

## Data Availability

Underlying raw data will be made available by the corresponding author upon reasonable request.

